# Decoding time-resolved neural representations of orientation ensemble perception

**DOI:** 10.1101/2023.09.29.560090

**Authors:** Ryuto Yashiro, Masataka Sawayama, Kaoru Amano

## Abstract

The visual system is capable of computing summary statistics of multiple visual elements at a glance. While numerous studies have demonstrated ensemble perception across different visual features, the timing at which the visual system forms an ensemble representation remains unclear. This is mainly because most previous studies did not uncover time-resolved neural representations during ensemble perception. Here we used orientation ensemble discrimination tasks along with EEG recordings to decode orientation representations over time while human observers discriminated an average of multiple orientations. We observed alternation in orientation representations over time, with stronger neural representations of the individual elements in a set of orientations, but we did not observe significantly strong representations of the average orientation at any time points. We also found that a cumulative average of the orientation representations over approximately 500 ms converged toward the average orientation. More importantly, this cumulative orientation representation significantly correlated with the individual difference in the perceived average orientation. These findings suggest that the visual system gradually extracts an orientation ensemble, which may be represented as a cumulative average of transient orientation signals, through selective processing of a subset of multiple orientations that occurs over several hundred milliseconds.

## Introduction

The visual system possesses remarkable abilities to extract the information required for specific situations among a plethora of inputs from the environment. Ensemble perception has been regarded as one such ability: the visual system is capable of extracting summary statistics of multiple elements (Whitney and Yamanashi Leib, 2018). Numerous studies have shown that ensemble perception can be formed in various dimensions from low- (Parkes et al., 2001; Bauer, 2009; Solomon, 2010) to high-level features (Haberman and Whitney, 2009; Yamanashi Leib et al., 2016). As the extraction of ensembles underlies various visual phenomena, including gist perception (Oliva and Torralba, 2006) and working memory (Brady and Alvarez, 2011; Lew and Vul, 2015), examining the characteristics of ensemble perception holds tremendous importance in the general understanding of visual processing.

Researchers have revealed several fundamental characteristics of ensemble perception. One prominent characteristic is rapid processing to extract the ensemble of multiple elements. Humans accurately judge the mean feature of multiple elements, even with a stimulus presentation of 50-100 ms (Chong and Treisman, 2003; Robitaille and Harris, 2011; Yamanashi Leib et al., 2016). In line with these studies, recent neuroimaging studies have successfully discriminated between two different stimulus groups by applying multivariate pattern analyses to EEG evoked signals around 100 ms post-stimulus (Roberts et al., 2019; Epstein and Emmanouil, 2021). Some studies have assumed parallel processing over the whole display as a mechanism of this rapid ensemble computation (Chong and Treisman, 2003; Robitaille and Harris, 2011; Corbett and Melcher, 2014; Cant et al., 2015).

On the other hand, an ensemble may be gradually computed through slower processing of multiple elements. Previous studies supported this idea by showing that the accuracy of ensemble discrimination improved with increasing stimulus duration (Li et al., 2016; Epstein et al., 2020). Slow processing lends itself to discounting the effect of outliers over time to achieve robust ensemble judgments (Haberman and Whitney, 2010; de Gardelle and Summerfield, 2011; Epstein et al., 2020), as the visual system is inherently sensitive to outliers in multiple objects (Cant and Xu, 2020). According to ideal observer models (Myczek and Simons, 2008; Solomon et al., 2011; Allik et al., 2013), this slow processing may be subserved by a serial and focused-attention mechanism over a subset of elements in the whole display (but see Solomon (2021)).

This controversy regarding the time course of ensemble perception has persisted for years, partly because most studies only used behavioral measures or neural signals with limited temporal resolution (Cant and Xu, 2012; Im et al., 2017; Cant and Xu, 2020; Tark et al., 2021), and consequently, the temporal dynamics of neural representation corresponding to ensemble perception remain unclear. Even with EEG recordings and time-resolved classification analyses (Roberts et al., 2019; Epstein and Emmanouil, 2021), precisely identifying the timing of ensemble perception is challenging because successful binary classification of different groups may potentially result from non-ensemble processing (e.g., the sampling of different individual elements within each group). To estimate precisely when ensemble perception was formed in the brain, we decoded the representational strength of both an ensemble and individual elements that constitute a stimulus set while human observers judged its average. Specifically, we conducted psychophysical experiments in combination with EEG recordings and time-resolved multivariate pattern analyses using orientation stimuli that are known to be decoded from EEG/MEG signals (Cichy et al., 2015; Pantazis et al., 2018; Hajonides et al., 2021). We also adopted a cross-decoding approach, in which we constructed orientation decoders within a single orientation discrimination task and applied these decoders to EEG signals during an orientation ensemble discrimination task. This approach allowed us to obtain the representational strength of multiple orientations in the form of probability scores. We found that different orientation representations exhibit alternating strength and are correlated with individual differences in the perceived average orientation when cumulatively averaged over approximately 500 ms. These results suggest that the visual system gradually computes an orientation ensemble over several hundred milliseconds.

## Materials and Methods

### Participants

Eleven paid volunteers (age: mean±s.d. = 25.0±3.84 years) participated in three experiments. All participants had normal or corrected-to-normal vision. All experiments were approved by the Ethics Committee of the University of Tokyo and were conducted in accordance with the Declaration of Helsinki guidelines. Written informed consent was obtained from all participants.

### Visual stimuli

Six Gabor patterns with a diameter of 4.8 deg (spatial frequency: 2.0 c/deg; Michelson contrast: 1) were presented against a gray background with a mean luminance of 62.5 cd/m2 (Fig. 1). The orientation of the Gabor patterns was determined according to the aim of each experiment, as explained in the following section, and each Gabor had a random phase in each trial. The center of the Gabor was located at a circle 5.8 deg from the fixation point. The stimulus was presented on a 24-inch LCD monitor (BenQ GW2480B) at a viewing distance of 100 cm.

**Figure 1.**
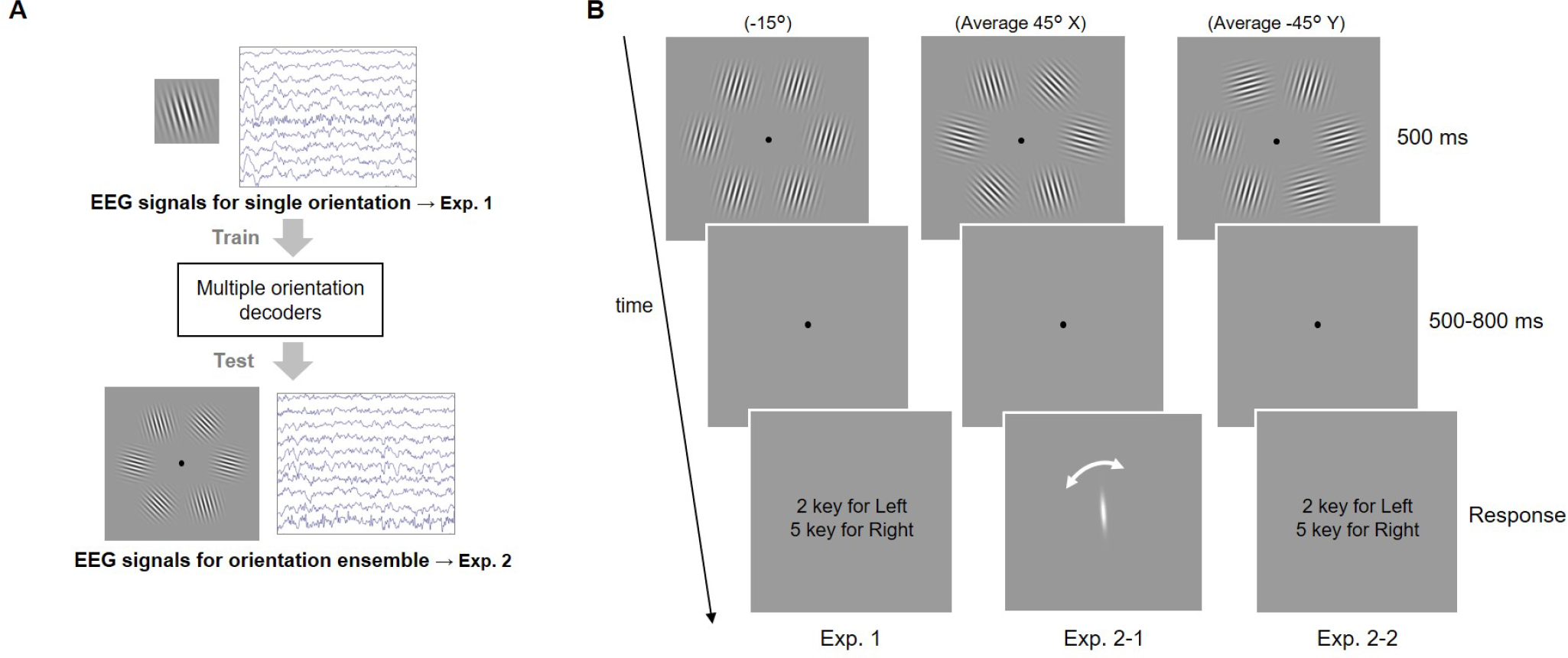
(A) Overview of experiments and analyses. We firstly recorded EEG responses to Gabor patterns consisting of uniform orientations (experiment 1) and used them to train orientation decoders that classify three different orientations (75°/45°/15° or -75°/-45°/-15°) for each time point. These decoders were then tested on the EEG signals that correspond to Gabor patterns with different orientations in the ensemble discrimination task (experiment 2). Then, we calculated three posterior probabilities that quantify the representational strength of three orientations during ensemble perception. (B) Schematic of psychophysical experiments. In all three experiments, we used six Gabor patterns presented in a circle. The task was to report the tilt of the patterns (Exp. 1) and the average orientation (Exp. 2-1 and 2-2). In Exp. 2-1, we directly measured the perceived average orientation using adjustment methods instead of the 2AFC task (Exp. 2-2).

### Experimental design

We conducted two types of experiments (experiments 1 and 2). In experiment 1, we conducted an EEG experiment with Gabor patterns with a single uniform orientation. Participants were presented with six Gabor patterns for 500 ms (Fig. 1B left, Exp.1). After a blank period of 500-800 ms, they were asked to judge if the patterns were tilted clockwise or counterclockwise relative to the vertical by pressing one of two keys. The next trial started 1500 ms after the response. All six patterns had identical orientation, which was randomly determined in each trial from six orientations ranging from -75° to 75° in steps of 30°, with 0° corresponding to the vertical. The response keys were randomly switched across blocks. Participants completed at least 6 blocks of 120 trials (each orientation was randomly presented 120 times across 6 blocks).

Experiment 2 consisted of behavioral and EEG experiments (experiments 2-1 and 2-2, respectively) with Gabor patterns with multiple orientations. In both experiments 2-1 and 2-2, participants were presented with a stimulus set of six Gabor patterns with different orientations (Fig. 1B middle and right, Table. 1 for four stimulus sets used in the experiment). Two of them had an average orientation of 45° and the others -45°. Because we were interested in how the decoding of the average orientation (45° or -45°) is affected by the presence of 45° or -45° patterns in the set, we created stimulus sets with and without elements corresponding to the average (X and Y, respectively) for each average orientation. In each trial, one of the sets was randomly presented to participants for 500 ms. The location of each element in the set randomly varied across trials.

In experiment 2-1, participants were asked to report the perceived average orientation by adjusting a white rotating bar presented at the center of the monitor (Fig. 1B middle) instead of a binary response. Note that we instructed participants to accurately report the average orientation they firstly perceived and not to postdictively change their response while rotating the bar. Also, we used four additional stimulus sets that served as dummy stimuli as well as the ones listed in Table 1 to prevent participants from being aware that the true average orientation was always 45° or -45° and reporting almost the same orientation across trials. The components of the dummy stimuli were as follows: (30°, 30°, 45°, 45°, 60°, 60°), (5°, 5°, 35°, 35°, 65°, 65°), (−30°, -30°, -45°, -45°, -60°, -60°) and (−5°, -5°, -35°, -35°, -65°, -65°). The trials in which the dummy stimuli were presented and the sign of the response was opposite to that of the true average (e.g., a trial with the response of -20° when the set had an average of 45°) were excluded in the following analyses.

**Table 1.** Four stimulus sets used in the experiment 2. The key difference between X and Y is whether the set contained the average orientation itself or not.

| Name | Components |
| --- | --- |
| Average 45° X | 15°, 15°, 45°, 45°, 75°, 75° |
| Average 45° Y | 15°, 15°, 15°, 75°, 75°, 75° |
| Average -45° X | -15°, -15°, -45°, -45°, -75°, -75° |
| Average -45° Y | -15°, -15°, -15°, -75°, -75°, -75° |

In experiment 2-2, the participants’ task was to indicate with a button press if the average orientation of the set was tilted clockwise or counterclockwise relative to the vertical (Fig. 1B right). The dummy stimuli used in experiments 2-1 were not presented. Participants completed at least 2 blocks of 120 trials in experiments 2-1 and 2-2 (each set was randomly presented 30 times in experiment 2-1 and 60 times in experiment 2-2). Note that in both experiments 1 and 2, we instructed participants not to attend to only a subset of the six patterns but to the whole display to ensure that participants did not explicitly change their strategy depending on the task demand, thereby enhancing the validity of our cross-decoding analysis (see below).

### EEG recording and preprocessing

In experiments 1 and 2-2, EEG signals were recorded with a sampling rate of 1000 Hz from 32 electrodes (BrainVision Recorder, BrainAmp amplifier, EasyCap; BrainProducts) which were located at FP1, FP2, F3, F4, F7, F8, Fz, T7, T8, C3, C4, Cz, FC1, FC2, FC5, FC6, P3, P4, P7, P8, Pz, TP9, TP10, CP1, CP2, CP5, CP6, PO3, PO4, O1, O2, and Oz. An electrode located between Fz and Cz served as a reference. Impedance for all the electrodes was kept below 5 kΩ throughout the experiments.

We preprocessed raw EEG signals using the EEGLAB toolbox for MATLAB. Raw EEG signals were downsampled to 500 Hz, band-pass filtered between 1 and 80 Hz, and epoched from 1000 ms before to 1500 ms after stimulus onset. We visually inspected the epochs to remove trials containing transient muscular activity and electrodes containing sustained noise. Epochs with incorrect responses were also removed. We then performed independent component analysis (ICA) and used the ICLabel plugin (Pion-Tonachini et al., 2019) to automatically obtain estimated labels for each component. Components with an estimated probability of more than 50% being artifact (i.e., a label of eye, muscle, heart, line noise, or channel noise) were removed, which resulted in 21±2.5 (mean±SD) components on average across participants and experiments. Epochs were baseline corrected relative to the prestimulus period of -300 to 0 ms. This procedure resulted in 922±136 (mean±SD, Exp.1) and 444±150 (mean±SD, Exp. 2-2) trials for further analyses.

### Orientation decoding

We constructed orientation decoders in a time-resolved manner for each participant, using potentials within a 100 ms time window and from 10 occipital and parietal electrodes (P3, P4, P7, P8, Pz, PO3, PO4, O1, O2, Oz) that were recorded in experiment 1 (Fig. 2) (cf. we used all 32 electrodes to construct decoders for our additional analyses, as shown in Discussion and Supplementary Figs. 3, 4, and 5). We trained two linear SVM classifiers (with a regularization parameter C of 1) for each participant to discriminate between three orientations included in each stimulus set of experiment 2 (i.e., 75°/45°/15° classifier or -75°/-45°/-15° classifier). Specifically, we firstly equalized the number of trials for each orientation in experiment 1 through undersampling. The EEG spatiotemporal patterns for each orientation were then divided into 5 sets, and the patterns within each set were averaged for each orientation to increase the signal-to-noise ratio and consequently decoding accuracy (Grootswagers et al., 2017). Four of the sets were used as training data and the remaining set as test data. The averaged signal within each set was z-scored using the mean and standard deviation of the training data over -300 ms to 900 ms relative to stimulus onset, in accordance with the procedure adopted in a previous study (Hermann et al., 2022). Subsequently, a linear multiclass SVM classifier was trained on the training data (12 samples = 4 sets × 3 orientations) and tested on the remaining 3 samples (1 set × 3 orientations). This procedure was repeated 5 times until all sets had served as test data (5-fold cross-validation). The whole procedure was repeated 20 times with random assignment of the patterns to each set. This resulted in 100 decoding accuracies, which were averaged together to obtain a single decoding accuracy for one time window. The 100 ms time window was shifted by a time step of 2 ms, and the same training and testing procedure was conducted, yielding time courses of 3-class orientation decoding accuracy over -300 ms to 900 ms relative to stimulus onset (Fig. 2 top right).

**Figure 2.**
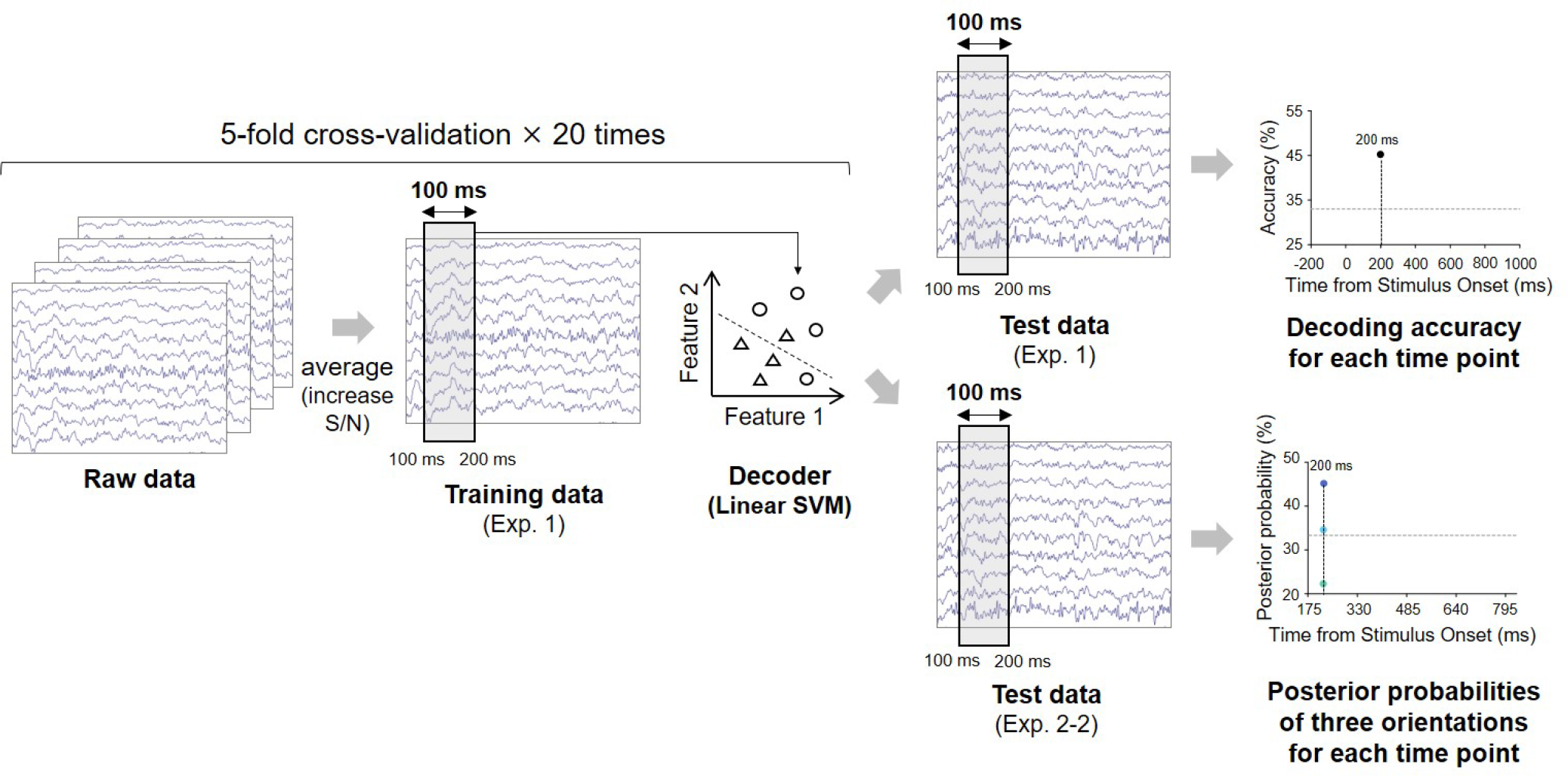
Procedure for the decoding analyses. We split the raw EEG data from Exp. 1 into five sets for each orientation (i.e., 75°, 45°, 15° or -75°, -45°, -15°) and averaged the data within each set (up to 40 trials) to increase the signal-to-noise ratio. Using the four averaged EEG patterns within a 100 ms time window for each orientation, we trained a linear SVM classifier, which was tested on the held-out three averaged EEG patterns (one for each orientation) from Exp. 1 to obtain an accuracy for the 3-class orientation decoding. We constructed multiple decoders independently for each time window. The time course of decoding accuracy was obtained after this procedure was repeated for the five cross-validation splits and with 20 different assignments of the raw EEG data to the five sets (top right). We also obtained the time courses of posterior probabilities of multiple orientations by testing the decoders on the EEG data from Exp. 2 (bottom right). All these analyses were performed individually for each participant.

### Decoding transient orientation representation during ensemble perception

We conducted a cross-decoding analysis to track how orientation representation fluctuates while participants performed an ensemble perception task. This analysis involved testing the classifier for one time window in experiment 1 on the temporally corresponding EEG signals from experiment 2 (Fig. 2 bottom right). Note that we used the EEG signals from experiment 2-2 that were averaged across 30 trials (almost equal to the number of trials included in each of the five sets when training the decoders) and z-scored as above. Then, we obtained three posterior probabilities of orientations (75°/45°/15° or -75°/-45°/-15°), which can be considered as the transient representational strength of each orientation (similar methods were employed by previous research (Rich and Wallis, 2016)). We defined SVM decision function values (obtained from sklearn.svm.LinearSVC.decision_function) that were passed through a softmax function as posterior probabilities for the test data. This procedure was conducted 100 times (5-fold cross-validation × 20 times) to generate 300 (= 3 orientations × 100 repetitions) probabilities in total, which were averaged across 100 repetitions to obtain three posterior probabilities for each orientation. We obtained time courses of posterior probabilities for three orientations by sliding the time window and repeating the same procedure.

### Estimating the timing of orientation ensemble representation formation

As shown in Results (Fig. 5), high posterior probabilities of 45° and -45° (expected results given the emergence of ensemble representation) were not observed for all four stimulus sets, suggesting that the neural representations corresponding to the ensemble orientation do not explicitly exist at specific time points. This led us to build a simple model to estimate how orientation representation evolved into ensemble orientation representation over time based on two assumptions: (1) an orientation whose posterior probability was maximum for one time point can be considered as most dominant in the orientation representation at that time point and (2) only the most dominant orientations are temporally averaged to form ensemble orientation representation in the brain (i.e., winner-take-all). According to these assumptions, we defined cumulative orientation representation as follows:

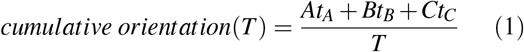

where *T* is elapsed time, (*A, B, C*) corresponds to (75°, 45°, 15°) and (−75°, -45°, -15°) for the stimulus set with an average orientation of 45° and -45°, respectively, and *t*_*A*_, *t*_*B*_, *t*_*C*_ denote how long each orientation had maximum posterior probability over *T* ms (Fig. 6). This value reflects the cumulative nature of ensemble processing, as suggested by previous studies (Haberman and Whitney, 2012; Epstein et al., 2020). This modeling procedure was conducted individually for each participant and stimulus set, thereby allowing us to estimate the dynamics of ensemble perception in the brain at the single-participant level.

Because perceived average orientation measured in experiment 2-1 differed across participants, we tested whether the estimated cumulative orientation representations significantly correlated with the perceived average orientations. Higher correlation at one time point makes it more likely that ensemble perception has been formed in the brain at that time point. Note that the correlations were calculated for the stimulus sets X and Y by collapsing the data across the true average orientation (45° or -45°). Specifically, we subtracted the true average orientation both from the estimated ensemble orientation representation and from the perceived average orientation. Then, we calculated the correlation between these two values for each time point, resulting in two time courses of the correlation for the stimulus sets X and Y.

### Statistical tests

We assessed the statistical significance of the orientation decoding accuracy using a cluster-based permutation test. We randomly permuted orientation labels for EEG data and recomputed orientation decoding accuracy according to the procedure described above. After repeating this procedure 1000 times, we generated null distributions of decoding accuracy for each time window, from which p-values for the real decoding accuracy were calculated. The original decoding accuracy curve was thus converted into a time course of p-values. Clusters were defined as neighboring time points with all p-values less than a significance level, and they were deemed significant if their size exceeded the threshold calculated from a null distribution of cluster size. Using the same procedure, we also assessed the statistical significance of the posterior probabilities.

Regarding correlation coefficients between the perceived average orientation and estimated cumulative orientation representation (calculated from Equation. 1), we again conducted a permutation test, obtaining null distributions of the correlation coefficients that were generated from 1000 repetitions of the above procedure for each time point. We assessed the statistical significance of the observed correlation coefficients for each stimulus set by comparing them with the 95th percentile of the null distribution.

## Results

We primarily sought to decode the dynamics in the representational strength of multiple orientations while human observers judged their average orientation. As mentioned in the Introduction, this investigation aimed to estimate the timing of ensemble representation formation in the brain. To this end, we conducted EEG recordings and psychophysical experiments. Figures 1 and 2 show an overview of our experiments and analyses. Eleven human adults participated in all experiments in which we used six Gabor patterns around a fixation point presented for 500 ms. We recorded EEG signals while participants discriminated Gabor patterns consisting of uniform orientations (experiment 1), and we independently constructed multiple classifiers (decoders) that could decode orientation from EEG signals at each time point (Fig. 1A and Fig. 2 top right). These decoders were used to investigate how orientation representations fluctuate over time while they engaged in the ensemble discrimination task (experiment 2). Specifically, these decoders were tested on the EEG signals recorded in Exp. 2 to obtain posterior probabilities of the three orientations that quantify the representational strength of these orientations (Fig. 1A and Fig. 2 bottom right).

### Orientation decoding

We first tested whether the uniform orientation of six Gabor patterns could be reliably decoded from EEG signals. Specifically, we trained and tested orientation decoders using EEG signals (from 10 occipital and parietal electrodes) evoked by six Gabor patterns with identical orientations (Fig. 1B left). Participants successfully discriminated the orientation of the patterns (tilted clockwise vs. counterclockwise relative to the vertical) in most trials with a mean accuracy of 98.4% (SD = 0.02%). An orientation decoder was constructed using EEG signals within a 100 ms time window that was shifted in steps of 2 ms, resulting in a time course of orientation decoding accuracy (Fig. 2 top right). Note that we constructed two types of decoders, each discriminating orientations contained in each stimulus set used in experiment 2 (i.e., 75°/45°/15° decoders and -75°/-45°/-15° decoders). Figure 3A shows that decoding accuracy reached significance at 174 ms and remained above chance level until 828 ms after stimulus onset (cluster-based permutation test; *p* < 0.05 cluster-defining threshold; *p* < 0.05 cluster-threshold). Only the decoders within this time range were used in the following analyses.

**Figure 3.**
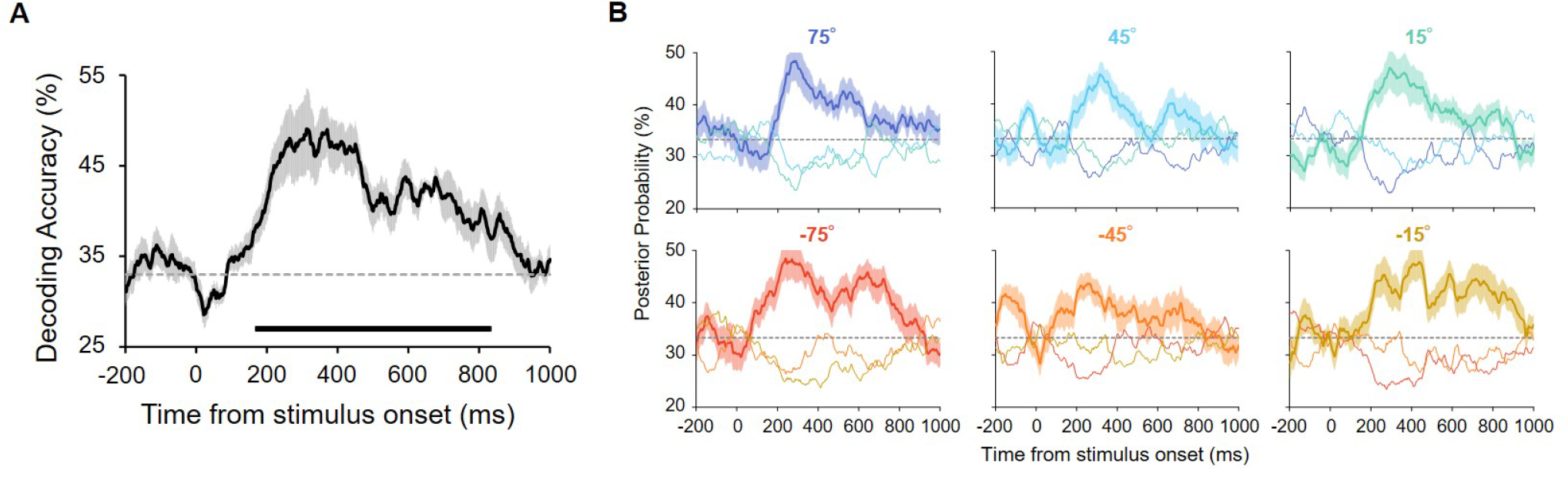
(A) Orientation decoding accuracy. Orientation was reliably decoded from 174 to 828 ms after stimulus onset. The chance level (33%) is shown as the dashed line. The shaded area indicates the standard error across participants. The horizontal line below the plot indicates time points that achieve significant decoding accuracy (revealed by a permutation test with cluster-defining threshold *p* < 0.05 and cluster threshold *p* < 0.05) (B) Posterior probabilities of presented orientations in Exp. 1. We computed these values by applying the decoders to the held-out test data from Exp. 1 and using the decoders’ output (see Methods for more details). Each panel shows the time course of the probability for each stimulus set with uniform orientations (shown above the panel). Solid lines correspond to results for the presented orientation. Probabilities for the other two orientations (e.g., 45° and 15° when 75° was presented) are shown as the thin lines. The probability of the presented orientation was higher for all the stimulus sets after stimulus onset, confirming that all of the physically presented orientations were reliably decoded and that the probability computed from the decoders’ output truly reflects the processing of the presented orientation.

To test whether all the six orientations were reliably decodable, we calculated the posterior probabilities of each orientation for each stimulus set by using the decoder’s output (see Methods). As shown in Fig. 3B, we obtained the highest probability of the presented orientation for each stimulus set throughout the stimulus presentation (e.g., the probability of 75° was higher than that of 45° and 15° when 75° was presented), ensuring that all the six orientations were reliably decoded. This result also indicates that the probabilities truly reflect the relative strength of orientation representations, an important implication for the analysis in the next section.

### Transient orientation representation during ensemble perception

The results of experiment 1 showed that orientation information was reliably decoded from EEG signals. Next, to quantify the strength of orientation representation during ensemble perception, we applied the decoders to the EEG signals in an orientation ensemble judgment task (experiment 2) within the corresponding time window and obtained the posterior probabilities of each orientation (Fig. 2 bottom right).

In Exp. 2, we used six Gabor patterns with two or three different orientations and an average of 45° or -45° (Table. 1). For each average orientation, we created two stimulus sets with and without elements corresponding to the average (X and Y) to see if the temporal dynamics of representational strength of each orientation are affected by the presence of the average orientation pattern in the set. We also recorded EEG signals while participants judged the average orientation of the set.

Then, the orientation decoders constructed in experiment 1 were tested on the EEG signals in experiment 2 to obtain the posterior probabilities of each orientation for each time point, which served as a measure of relative strength of the transient representations of each orientation during orientation ensemble perception (Fig. 2 bottom right), as confirmed in Exp. 1 (Fig. 3B). Based on these probabilities, we hypothesized the following: if ensemble representation is explicitly formed at a specific time point and in certain areas of the brain, as assumed in previous studies (Haberman and Whitney, 2012; Baek and Chong, 2020a; Utochkin et al., 2023), significantly high probability of the true average orientation (i.e., 45° or -45°) would be observed at that time point, allowing us to estimate the precise timing of ensemble perception formation.

This prediction regarding the emergence of ensemble representation is built on the assumption that participants successfully perceived the average orientation as close to the true average (45° or -45°). To directly assess the precision of the perceived average orientation of the six Gabor patterns, we initially performed a psychophysical experiment using adjustment methods (experiment 2-1; Fig. 1B middle). Participants reported the perceived average orientation on a continuous scale by rotating a bar stimulus. Figure 4 shows the perceived average orientations for each participant and stimulus set. Although the perceived orientations varied across participants, they were mostly close to the true average, indicating that participants perceived the average orientation fairly well.

**Figure 4.**
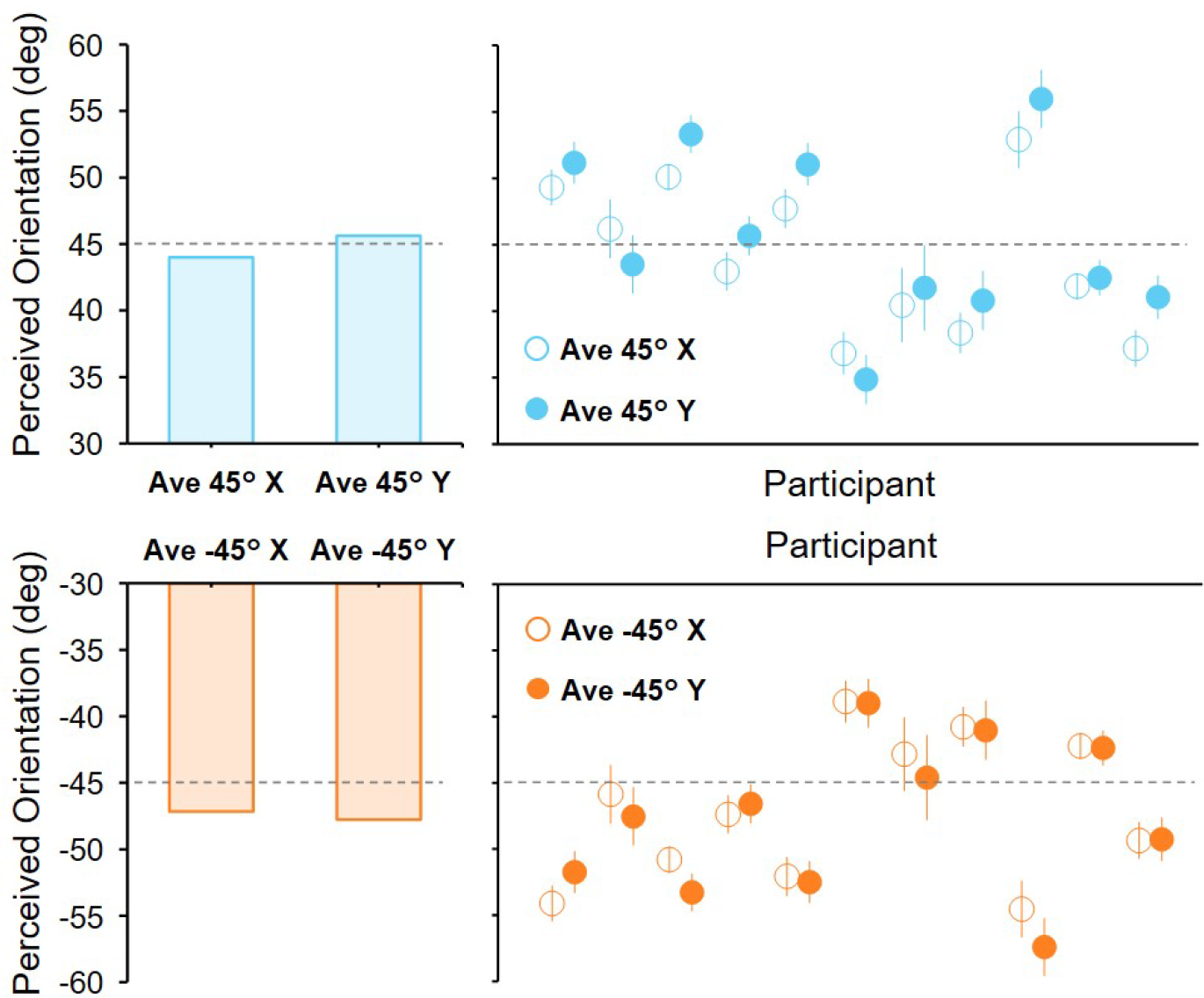
Perceived average orientations in Exp. 2-1. The left bar plots show the perceived orientation averaged across participants. Each circle in the right panels represents the perceived orientation for each participant. Error bars indicate the standard error across trials. The true average is shown as a dashed line. Most participants accurately perceived the average orientation, with errors mostly within a range of ±10°.

**Figure 5.**
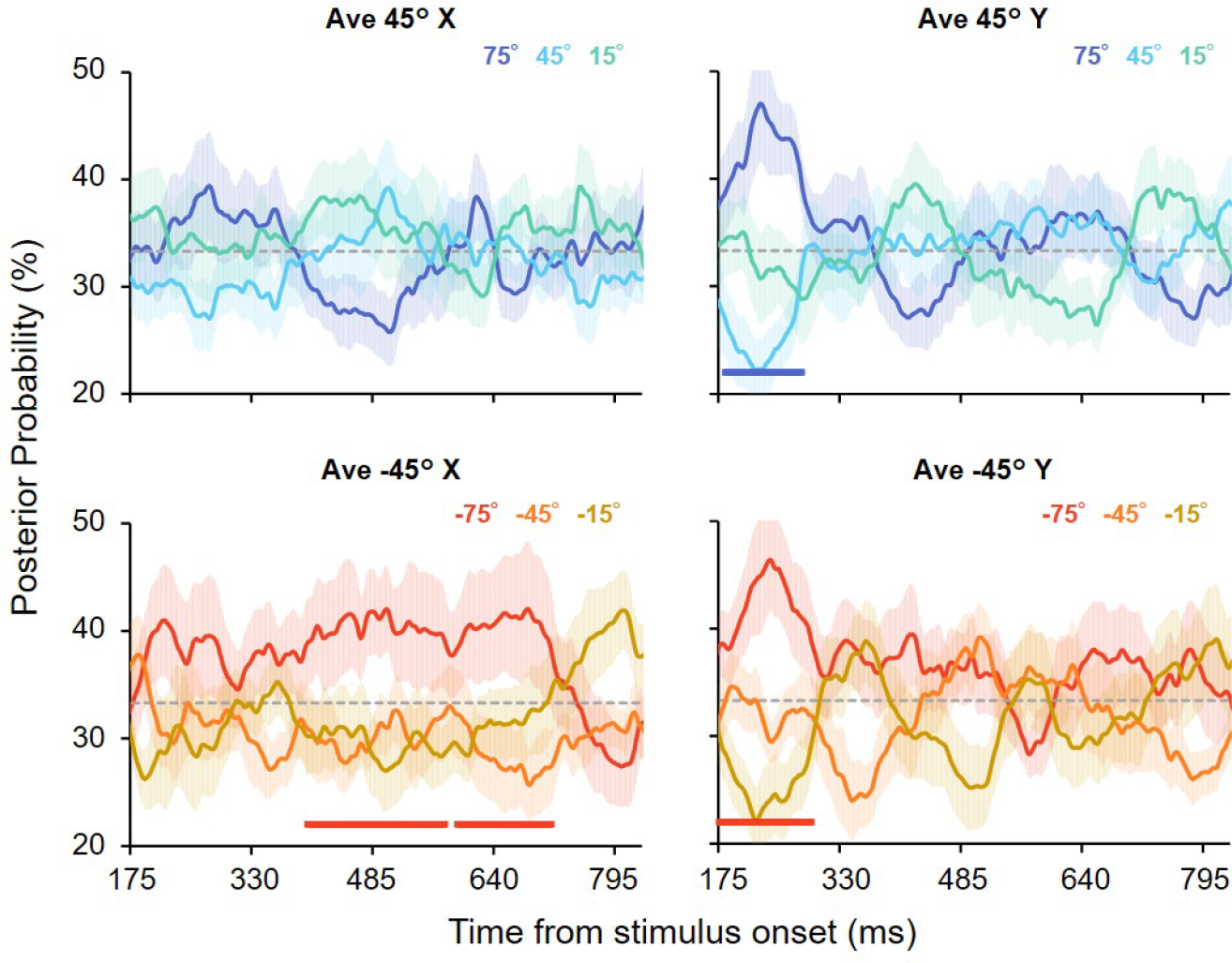
Posterior probabilities of orientations that constituted the stimulus sets in Exp. 2. Each panel shows results for each stimulus set. Each color corresponds to a specific orientation. The time range is restricted to the period where above-chance decoding accuracy was achieved (Fig. 3A). The shaded area shows the standard error across participants. The probabilities of the true average (45° and -45°, shown as the light blue and orange lines, respectively) were not high across all the time points. Also, significantly high probabilities of individual orientations (75° or -75°) were observed, indicated by the horizontal lines below (permutation tests with cluster-defining threshold *p* < 0.1 ; cluster threshold *p* < 0.1).

**Figure 6.**
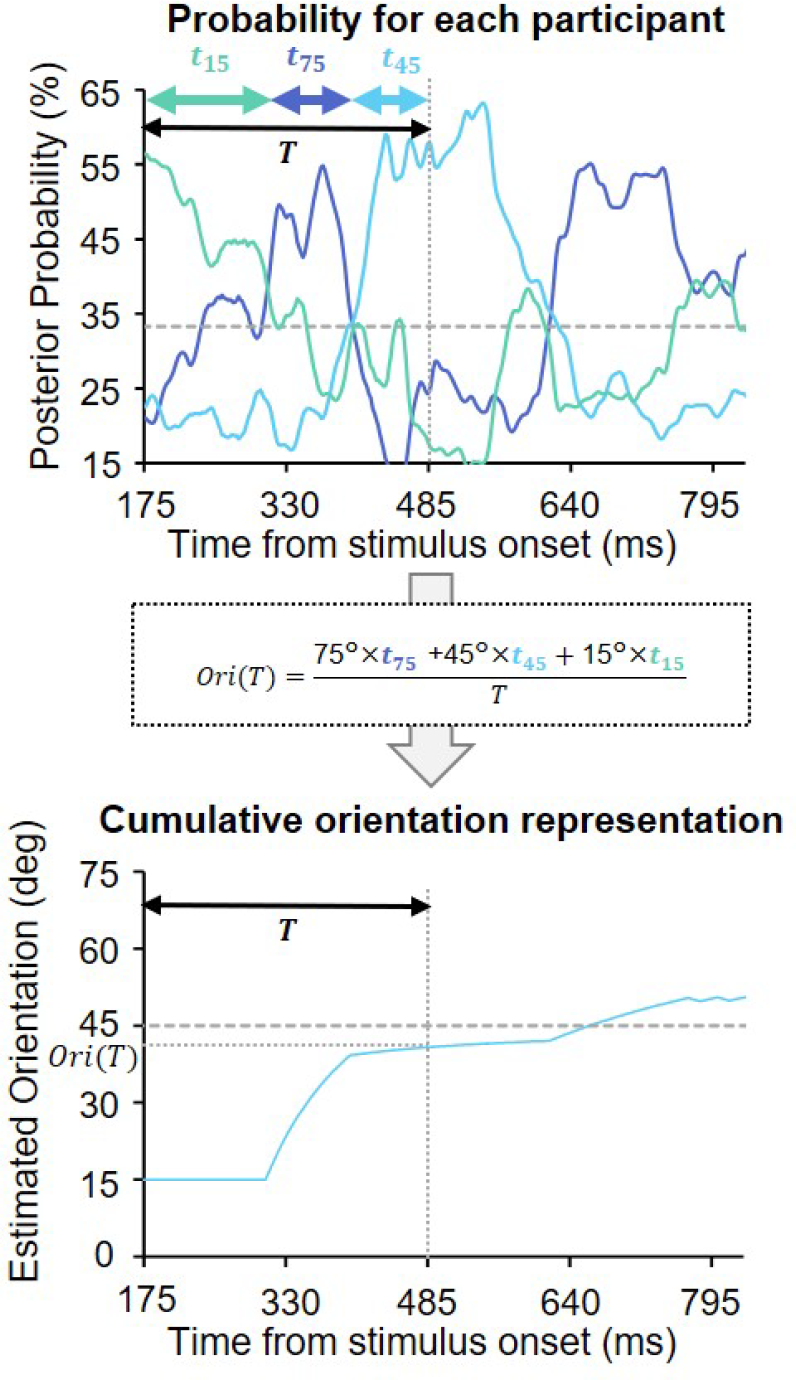
Procedure for estimating the cumulative orientation representation over time. The time courses of the posterior probabilities were used as transient orientation representations. The top panel shows the posterior probabilities for one participant. To estimate the cumulative orientation representation value at one time point for this participant, we counted the number of time points at which each orientation had the highest probability up until that time point. These counts were then used to compute a weighted average of the three orientations, which corresponds to the estimated orientation representation for that time point. This procedure was repeated for all time points to uncover how orientation representation evolves over time.

We then conducted experiment 2-2 in which EEG signals were recorded while participants discriminated the average orientation of the set (2AFC discrimination task, Fig. 1B right; behavioral performance was 92.6 % on average (SD = 0.10%)). As described above, we obtained the time courses of the posterior probabilities of three orientations for each stimulus set (Fig. 5). Contrary to our initial hypothesis, for set Y (without 45° or -45° patterns), the probabilities of the true average were not significantly higher than those of the other orientations throughout the entire period. Notably, the same held true even for set X that contained the true average orientation as a local element. We obtained qualitatively similar results even if we constructed decoders using all 32 electrodes (including frontal and central ones as well as the original 10 occipital and parietal electrodes) (Supplementary Fig. 5). These results suggest that ensemble orientation may not be explicitly represented at a specific time point and in certain areas of the brain, irrespective of whether the set contains the average orientation itself as a local element.

Although the probabilities mostly hovered near the chance level (33%) due to the great individual differences in temporal profiles of the probability across participants (Suppl Fig. 1), we observed significantly higher probabilities of 75°/-75° at the group level, which lasted until around 300 ms post-stimulus for two of the four stimulus sets (average 45° Y and average -45° Y) and in the later period for one stimulus set (average -45° X; cluster-based permutation test; *p* < 0.1 cluster-defining threshold; *p* < 0.1 cluster-threshold). Given that 75° and -75° are the most informative for the task goal, the visual system may selectively process task-relevant information, especially during an early stage of ensemble processing.

### Ensemble representation as a cumulative average of transient representations

As we did not explicitly observe high posterior probability of the true average, we next hypothesized that ensemble representation would emerge not as a distinct signal at specific time points but as a cumulative average of the transient orientation representations. This idea of cumulative ensemble computation was partly inspired by the iterative process for ensemble perception proposed in previous studies (Haberman and Whitney, 2012; Epstein et al., 2020), in which ensemble representation was assumed to gradually be formed over time through feedforward and feedback processing between visual areas. Thus, we next sought to correlate the perceived average orientation for each participant (Exp. 2-1) with a cumulative average of the orientation representations (Exp. 2-2) at each time point to estimate when ensemble perception was formed in the brain. We expected that if ensemble representation was cumulatively formed from stimulus onset until a specific time point, a significantly high correlation would be observed at that time point.

Figure 6 shows the procedure for estimating the cumulative orientation representation. For each time point, we computed a cumulative average of three orientations (75°/45°/15° or -75°/-45°/-15°) that exhibited the maximum probability up to that time point (see Methods for more details). This was repeated for all time points to obtain a time course of cumulative orientation representation for each participant and stimulus set. The time course differed considerably across participants, while cumulative orientation representation mostly converged close to the average orientation, (Supplmentary Fig. 2) indicating that orientation representation converges to the final ensemble representation through distinct dynamics for each participant.

Then, we computed the correlation between the cumulatively estimated orientation representations and perceived average orientations of each participant in Exp. 2-1 for the stimulus sets X and Y (the sign of the average orientation was ignored). Fig. 7 shows the correlation coefficient for each time point. We observed a gradual increase in the correlation coefficients between the neurally estimated cumulative orientation representation and perceived average orientation over time. Permutation tests with null distributions of the correlation coefficient revealed that significantly high correlations persisted from 420 ms (set X) and 518 ms (set Y) after stimulus onset. These results suggest that ensemble perception can be formed as a temporal average of the transient orientation representations over approximately 500 ms after stimulus onset through distinct temporal evolution of orientation representation across individuals.

**Figure 7.**
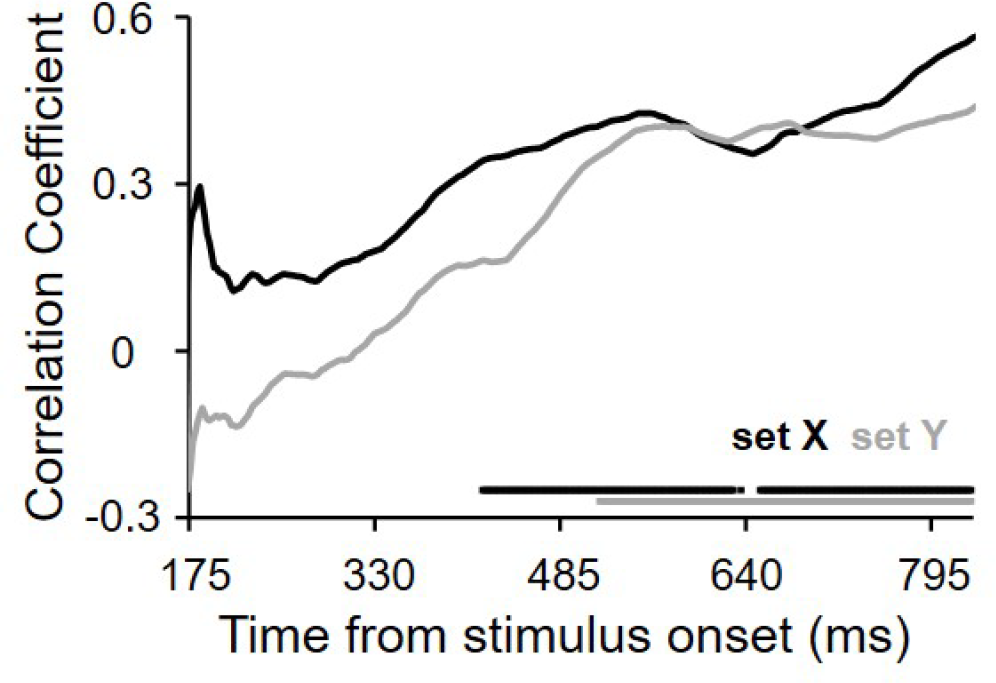
Correlations between the perceived average orientation and orientation representation estimated from the EEG signals (Fig. 6) for the two set types. They gradually increased over time and reached significance at 420 ms and 518 ms after stimulus onset for sets X and Y, respectively. The horizontal lines below show time points at which correlation was statistically significant (permutation tests, *p* < 0.05).

## Discussion

We uncovered the temporal dynamics of orientation ensemble perception by employing time-resolved multivariate pattern analyses on EEG signals and psychophysical experiments. One of the keys to achieve this was the use of cross-decoding analysis for each time point between the experiments (i.e., single and ensemble orientation discrimination task; Exp. 1 and 2-2, Fig. 1) rather than simple binary classification of two different stimulus sets within an experiment (Roberts et al., 2019; Epstein and Emmanouil, 2021). Another key is that the neural orientation representations predicted from the temporally averaged output of multiple orientation decoders were directly correlated with the perceived average orientations in the task with adjustment methods. The time course of the correlation enabled us to precisely estimate when ensemble perception is formed in the brain, which would have otherwise remained elusive with behavioral paradigms using discrete responses that have been adopted in most studies on ensemble perception (i.e., member identification task or 2AFC task).

One notable finding is that significantly strong orientation representation corresponding to the true average (45° or -45°) did not appear at any time points across participants (Fig. 5). This contrasts with our initial prediction that regardless of the presence of the 45° and -45° patterns in the set, the neural representation of the true average orientation will emerge when ensemble representation is formed in the brain (this prediction was partly motivated by the recent finding of shared neural representations between the perception of physical stimuli and mental imagery (Fukuma et al., 2022)). In addition, we found that a cumulative average of the output of the decoders (i.e., transient orientation representations) over time is significantly correlated with the perceived average orientation (Fig. 6 and Fig. 7). Taken together, although existing computational models (Haberman and Whitney, 2012; Baek and Chong, 2020a; Utochkin et al., 2023) and fMRI studies (Cant and Xu, 2012; Tark et al., 2021) indicated the presence of ensemble representation in higher areas of the brain, an ensemble of multiple orientations may not be explicitly represented in specific areas but represented as a temporal summation of distributed response patterns of multiple orientation-selective neurons in lower visual areas. This possibility is further supported by our additional analyses in which we constructed orientation decoders using 32 EEG electrodes, including frontal ones (note that we used only 10 occipital and parietal electrodes in our original analyses) (Supplementary Fig. 5). With these decoders that potentially reflect neural signals in both lower and higher areas, we still did not observe high posterior probabilities of the true average.

We also found uneven representational strength of multiple orientations, as reflected in the higher posterior probability of 75° and -75° in Fig. 5. This suggests that the visual system may selectively process individual task-relevant elements, especially in an early stage of ensemble processing. The notion of selective processing is corroborated by a recent fMRI study showing a transient BOLD signal change in response to a few outliers in a collection of objects (Cant and Xu, 2020), and by behavioral studies showing that humans place unequal weights on visual elements distributed over space to make judgments on their ensemble (de Gardelle and Summerfield, 2011; Li et al., 2017; Lau and Brady, 2018; Pascucci et al., 2021). In line with these findings, our study provides complementary evidence for selective processing underlying ensemble perception in terms of the relative strength of neural representations of multiple elements.

The time courses of correlations between the participants’ perceived average orientation and cumulative average of neurally predicted orientation representation revealed that orientation ensemble perception occurs approximately 500 ms after stimulus onset (Fig. 7). However, previous behavioral studies have shown that discrimination performance was invariant to stimulus duration, suggesting a much shorter timescale for the computation of ensembles (Chong and Treisman, 2003; Robitaille and Harris, 2011; Yamanashi Leib et al., 2016). This is further supported by EEG studies that demonstrated reliable discrimination of two stimulus sets with different ensembles using early evoked EEG multivariate patterns (Roberts et al., 2019; Epstein and Emmanouil, 2021). How can we reconcile these results? One crucial problem with previous behavioral studies (Chong and Treisman, 2003; Robitaille and Harris, 2011) is that they used unmasked visual stimuli that allowed observers to access stimulus information beyond the actual stimulus duration, potentially influencing the behavioral performance. In fact, some studies indicated that backward masking degraded the performance of ensemble discrimination, especially with short stimulus duration (Whiting and Oriet, 2011; Epstein et al., 2020). Additionally, reaction times for ensemble judgments are typically within the range of 600 to 1000 ms (Robitaille and Harris, 2011; Li et al., 2016), which is consistent with our findings given the delay associated with motor response. Furthermore, in previous EEG studies, it is possible that distinct multivariate EEG patterns for the two stimulus sets could merely reflect the encoding of different individual elements within each set. Specifically, the processing of a subset of elements, rather than the formation of ensemble representation from different sets, would potentially yield significant classification accuracy of the two sets observed early in time. Considering these facts and possibilities, it would be reasonable to assume that the visual system gradually forms ensemble perception over several hundred milliseconds.

One might argue that our results are potentially influenced by the low (though statistically reliable) decoding performance (Fig. 3) and choice of classifiers. This is reasonable given that our main findings are all based on the output of the classifiers. To address this concern, we employed several strategies to maximize the signal-to-noise ratio of the EEG data and consequently improve decoding accuracy (Grootswagers et al., 2017). These strategies included averaging the EEG data across trials and using multiple data points within each time window as features for the classifier. We also analyzed the data using different classifiers (e.g., nonlinear SVM) but still obtained qualitatively similar results (Supplementary. Fig. 3 for decoding accuracy, Supplementary. Fig. 4 and 5 for posterior probability of each orientation). Therefore, we draw robust conclusions regarding the presence of explicit ensemble representation and the time course of orientation ensemble perception, minimizing potential biases associated with the decoding performance and classifier selection.

Overall, our results support slow ensemble computation with selective processing over a subset of multiple orientations. Aligning our results with the insights from previous studies (Myczek and Simons, 2008; Solomon et al., 2011; Allik et al., 2013), we infer computational mechanisms of ensemble perception: serial and focused-attention processing may underlie the slow and selective ensemble computation. However, this does not entirely preclude the involvement of parallel processing in computing an ensemble. A combination of parallel and serial processing may underpin ensemble perception (Whitney and Yamanashi Leib, 2018; Baek and Chong, 2020b). For instance, a few elements may be processed in parallel, with unequal weight assigned to each element, which can occur serially across different sets of elements over time. Alternatively, an individual element may be serially sampled, followed by a one-off ensemble computation instead of the cumulative ensemble computation we assumed. Further studies will be necessary to examine these possibilities and delineate the computational mechanisms of ensemble perception in more detail. Comparison of fMRI and EEG/MEG, along with a similarity analysis using dissimilarity matrices from the two modalities (Cichy et al., 2014; Hebart and Baker, 2018), could take advantage of both high spatial and temporal resolution, which is conducive to a deep understanding of how and when ensemble representations are formed in the brain. Also, as covert attention focused on specific elements or distributed over space is inherently involved in the extraction of ensembles (Chong and Treisman, 2005; Emmanouil and Treisman, 2008; Brand et al., 2012), decoding covertly attended visual stimuli (Noah et al., 2020) will facilitate further investigation of the temporal dynamics and computational mechanisms of ensemble perception.

## Acknowledgements

This study was supported by JST SPRING, Grant Number JPMJSP2108.

## Author contributions statement

R.Y., S.M., and K.A. conceived and designed the experiments. R.Y.. conducted the experiments and analyzed the data. R.Y., S.M., and K.A wrote the manuscript.

## Supplementary Figures

**Supplementary Figure 1.**
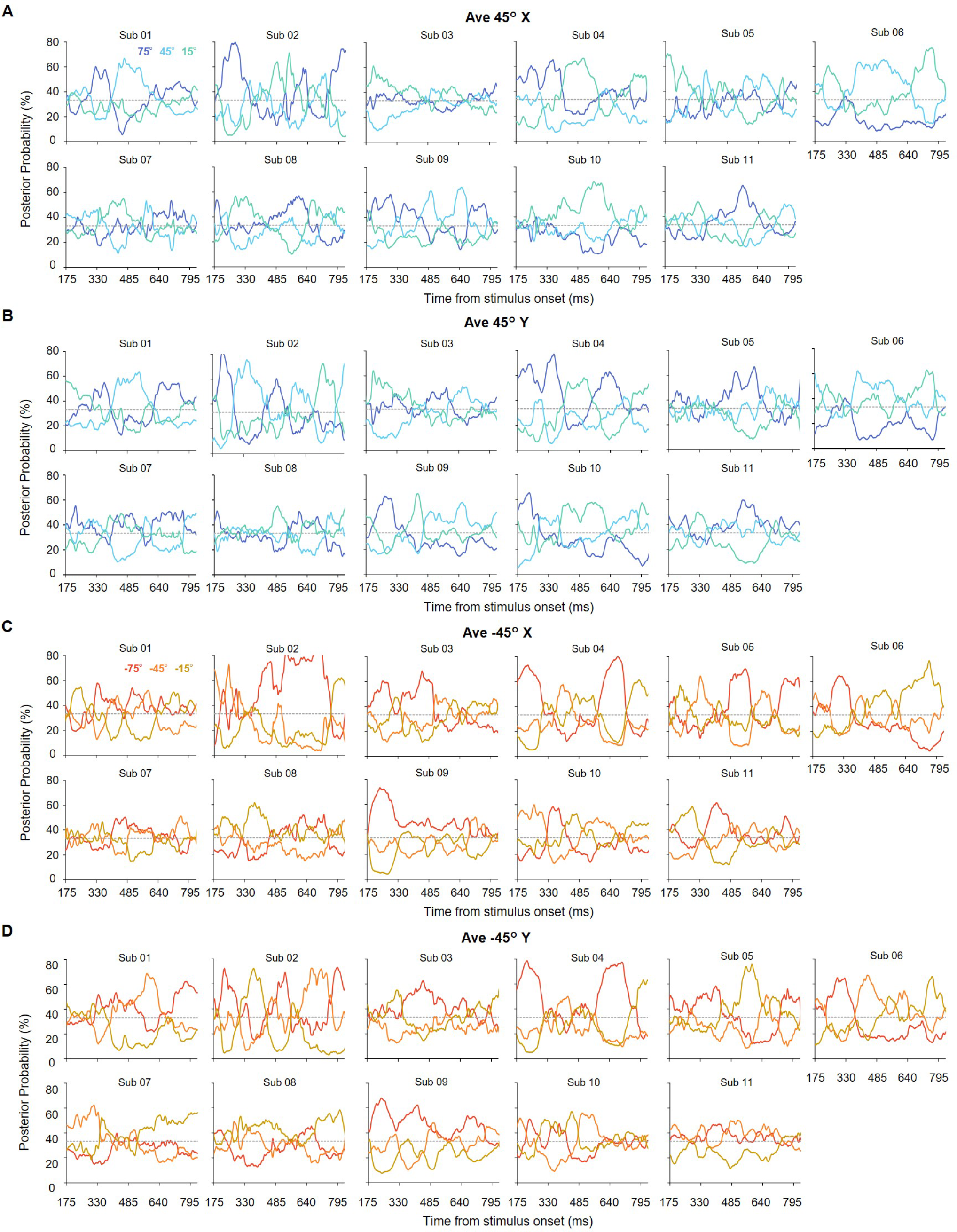
Time courses of posterior probabilities in Exp. 2 for 11 participants (sub 01-11). Each panel (A-D) shows the results for each stimulus set. We observed distinct temporal profiles of the probabilities (i.e., relative representational strength of multiple orientations) across participants. All other conventions are the same as those in Fig. 5.

**Supplementary Figure 2.**
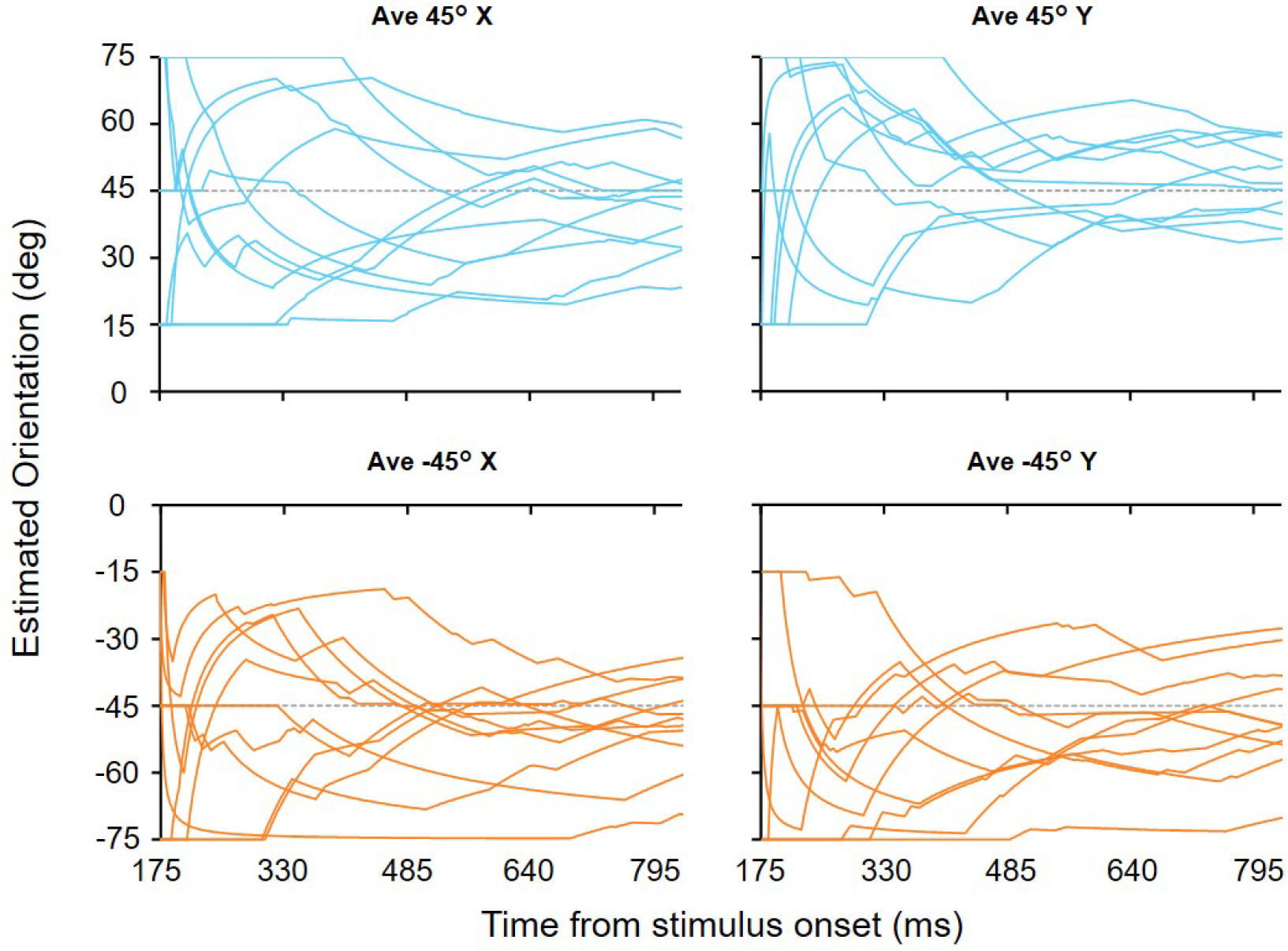
Fluctuations in the cumulative orientation representation for each stimulus set. Each line represents the result for each participant. The true averages are shown as dashed lines. Although estimated orientation representations exhibited distinct dynamics across participants, they mostly converged to a value close to the true average.

**Supplementary Figure 3.**
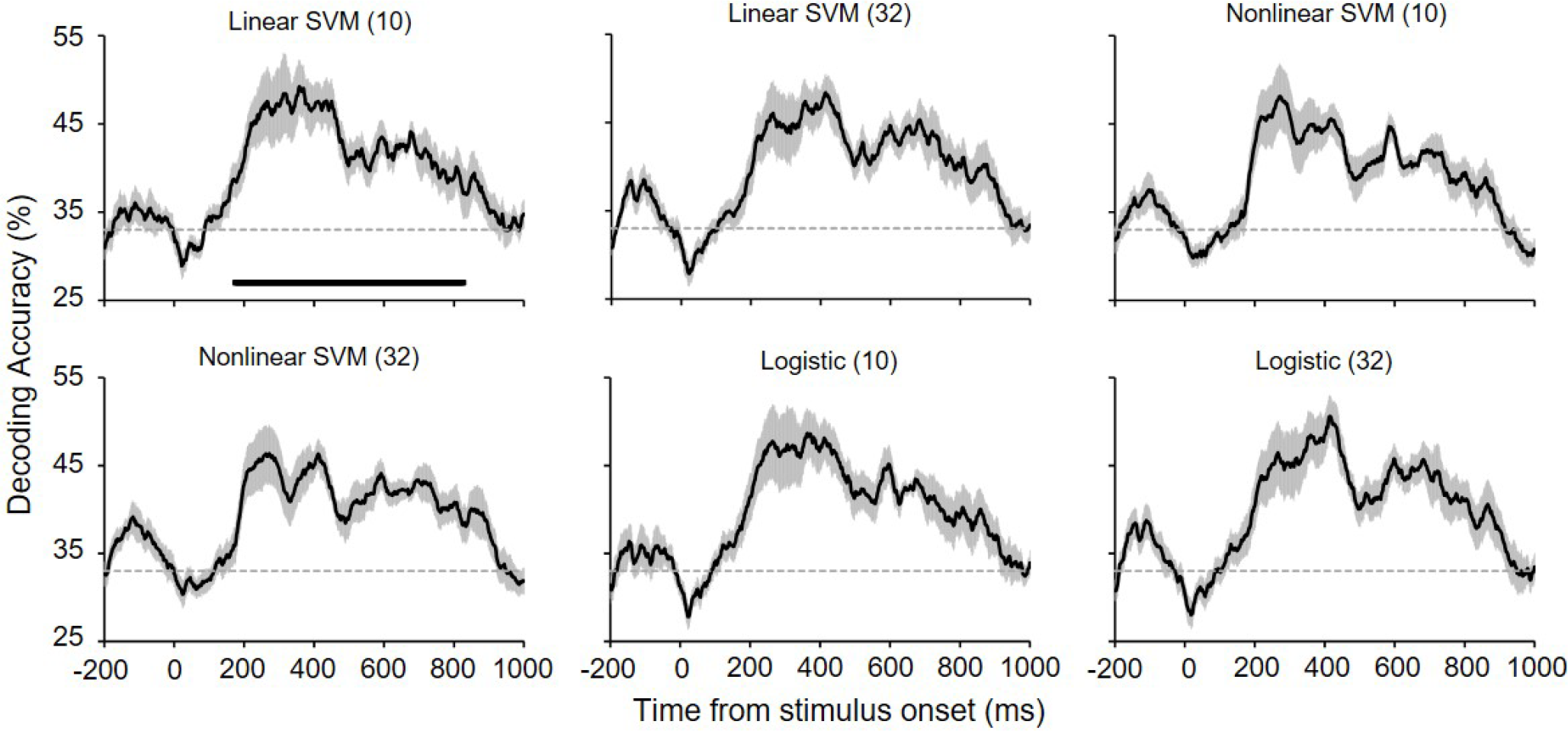
Orientation decoding accuracy for different decoder types (Linear SVM, Nonlinear SVM, or Logistic regression) and number of electrodes used to construct decoders (in brackets; 10 or 32). The top left panel is the same as Fig. 3A. The decoding accuracy profiles remained qualitatively similar, even when we used different types of decoders and all 32 electrodes. All other conventions are the same as in Fig. 3A.

**Supplementary Figure 4.**
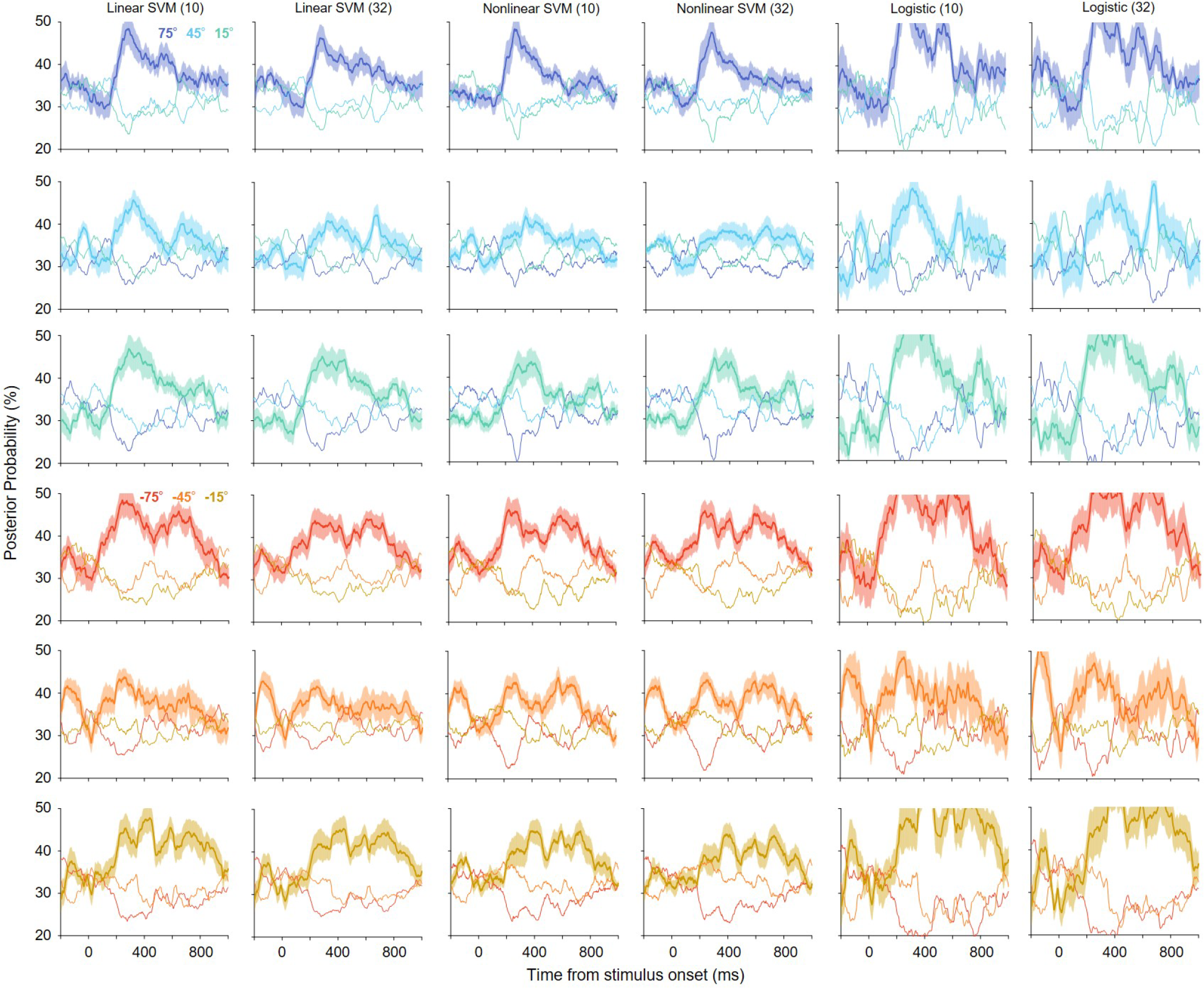
Posterior probabilities of six orientations in Exp. 1 for different decoder types and numbers of electrodes used to construct decoders (in brackets; 10 or 32). The panels in the leftmost column are the same as those in Fig. 3B. The probability profiles were qualitatively similar across different types of decoders and electrodes. All other conventions are the same as in Fig. 3B.

**Supplementary Figure 5.**
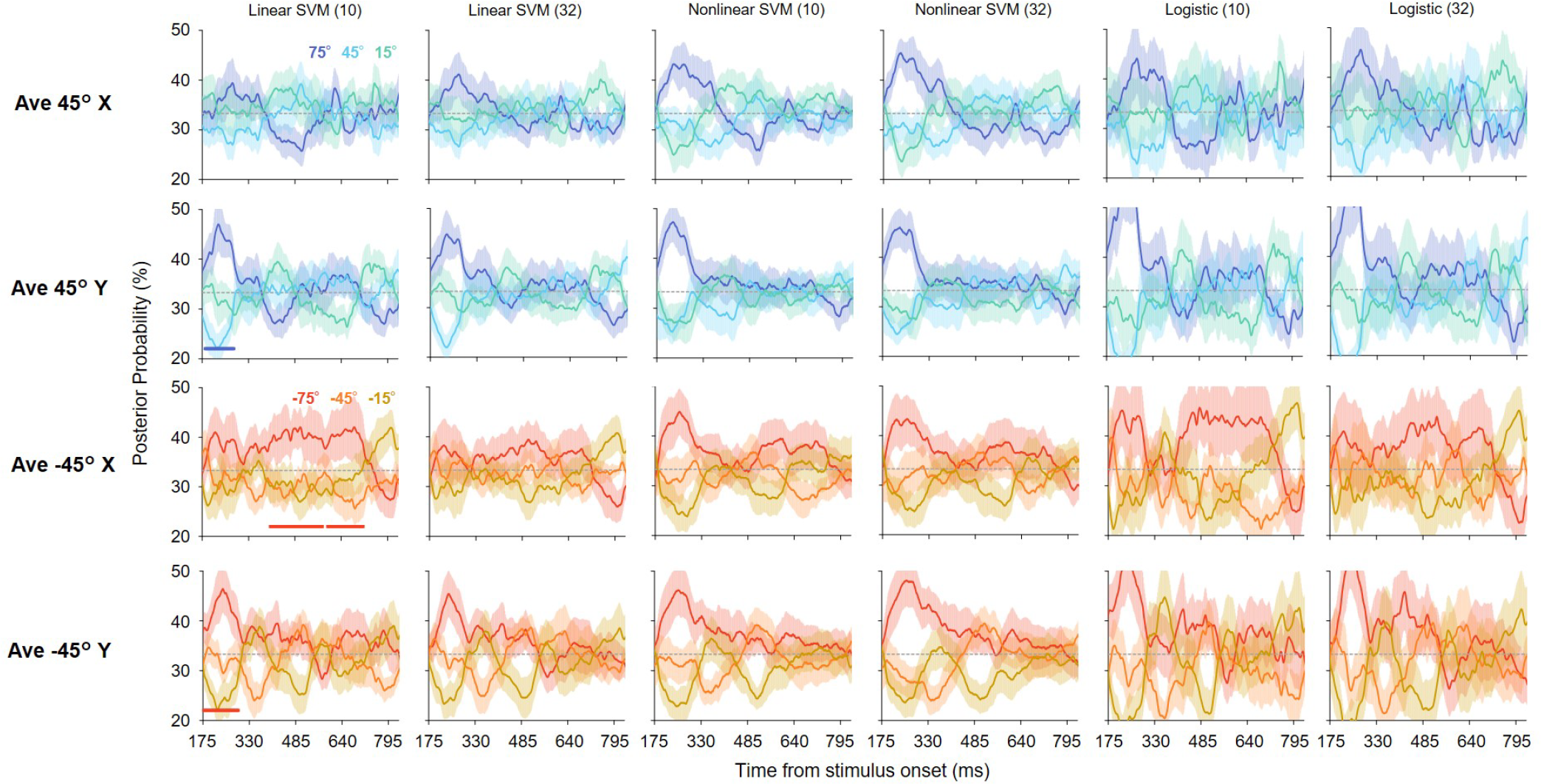
Time courses of posterior probabilities of orientations included in the stimulus sets in Exp. 2 for different decoder types and number of electrodes used to construct decoders (in brackets; 10 or 32) The panels in the leftmost column are the same as those in Fig. 5. The probability profiles were qualitatively similar across different types of decoders and electrodes. All other conventions are the same as in Fig. 5.

